# Tunturi virus isolates and metagenome-assembled viral genomes provide insights into the virome of *Acidobacteriota* in Arctic tundra soils

**DOI:** 10.1101/2024.08.08.607240

**Authors:** Tatiana Demina, Heli Marttila, Igor S. Pessi, Minna K. Männistö, Bas E. Dutilh, Simon Roux, Jenni Hultman

## Abstract

**Background:** Arctic soils are climate-critical areas, where microorganisms play crucial roles in nutrient cycling processes. *Acidobacteriota* are phylogenetically and physiologically diverse bacteria that are abundant and active in Arctic tundra soils. Still, surprisingly little is known about acidobacterial viruses in general and those residing in the Arctic in particular. Here, we applied both culture-dependent and -independent methods to study the virome of *Acidobacteriota* in Arctic soils.

**Results:** Five virus isolates, Tunturi 1-5, were obtained from Arctic tundra soils, Kilpisjärvi, Finland (69°N), using *Tunturibacter* spp. strains originating from the same area as hosts. The new virus isolates have tailed particles with podo-(Tunturi 1, 2, 3), sipho-(Tunturi 4), or myovirus-like (Tunturi 5) morphologies. The dsDNA genomes of the viral isolates are 63–98 kbp long, except Tunturi 5, which is a jumbo phage with a 309-kbp genome. Tunturi 1 and Tunturi 2 share 88% overall nucleotide identity, while the other three are not related to one another. For over half of the open reading frames in Tunturi genomes, no functions could be predicted. To further assess the *Acidobacteriota*-associated viral diversity in Kilpisjärvi soils, bulk metagenomes from the same soils were explored and a total of 1881 viral operational taxonomic units (vOTUs) were bioinformatically predicted. Almost all vOTUs (98%) were assigned to the class *Caudoviricetes*. For 125 vOTUs, including five (near-)complete ones, *Acidobacteriota* hosts were predicted. *Acidobacteriota*-linked vOTUs were abundant across sites, especially in fens. *Terriglobia*-associated proviruses were observed in Kilpisjärvi soils, being related to proviruses from distant soils and other biomes. Approximately genus- or higher-level similarities were found between Tunturi viruses, Kilpisjärvi vOTUs, and other soil vOTUs, suggesting some shared groups of *Acidobacteriota* viruses across soils.

**Conclusions:** This study provides acidobacterial virus isolates as laboratory models for future research and adds insights into the diversity of viral communities associated with *Acidobacteriota* in tundra soils. Predicted virus-host links and viral gene functions suggest various interactions between viruses and their host microorganisms. Largely unknown sequences in the isolates and metagenome-assembled viral genomes highlight a need for more extensive sampling of Arctic soils to better understand viral functions and contributions to ecosystem-wide cycling processes in the Arctic.

## Introduction

Huge amounts of carbon and nitrogen are accumulated in Arctic permafrost-affected tundra soils and are expected to be released with increasing temperatures due to climate change [1]. Despite seemingly harsh environmental conditions, Arctic soils host diverse and active microbial communities that decompose soil organic matter and contribute to greenhouse gas cycling [2–6]. Arctic soil microbial communities have been shown to experience compositional and functional changes with permafrost thawing [7–12]. However, the Arctic covers a large geographic area with diverse environments, and more studies are needed to understand and predict the responses of microorganisms to climate change, including the increased amounts of soil carbon and nitrogen being released from Arctic permafrost. In particular, the contribution of viruses to ecosystem-wide cycling processes in soils has been little explored compared to aquatic environments [13–15].

Soil viruses are abundant and diverse [13,16] and have various impacts on their host microorganisms [17,18]. By lysing microbial cells, viruses indirectly affect biogeochemical cycling in soil [19]. With lysogenic conversion, i.e., the expression of genes encoded by a lysogen, temperate viruses may support fitness of their hosts, e.g., by protecting them from other viral infections [20] or increasing their ability to form biofilms [21]. Moreover, a variety of auxiliary metabolic genes (AMGs) have been predicted in soil viruses [22] and some of soil virus AMGs involved in carbon cycling have been experimentally confirmed to be active [23,24]. Finally, experimental warming of tundra soils from the Alaskan permafrost region resulted in increased abundance of viral AMGs (glycoside hydrolases) in warmed soils [25].

*Acidobacteriota* are widespread and abundant in various environments, especially in acidic soils [26–30]. Despite their abundance, a relatively limited number of *Acidobacteriota* species have been isolated and characterized, as their cultivation is often challenging [31,32]. While information on their ecological functions remains fragmentary, soil *Acidobacteriota* seem to have significant roles in carbon [33–37], nitrogen [6] and sulfur [38] cycling. Moreover, *Acidobacteriota* include many putative keystone taxa, i.e., strong drivers of microbiome structure and functioning, in soils and as part of plant-associated microbiota [39]. *Acidobacteriota* are phylogenetically diverse and comprise 15 taxonomic classes [40]. The members of the class *Terriglobia* are typically dominant and active in Arctic tundra and boreal forest soils [6,28,41,42] and are represented by several cultured isolates [37,43–45]. Recently, the genus *Tunturibacter* has been described in the class *Terriglobia* [46]. The type species, *Tunturibacter lichenicola*, was originally known as *Edaphobacter lichenicola* [45]. *Tunturibacter* representatives are Gram-negative aerobic rods, produce extracellular polysaccharide-like substances and are able to hydrolyze various polysaccharides [46].

Surprisingly little is known about viruses that infect *Acidobacteriota*. With the use of genomic data, proviral sequences have been predicted in acidobacterial genomes [47], and viral populations extracted from metagenomic datasets have been putatively linked to acidobacterial hosts [23,25,48,49]. Here, we used both culture-dependent and -independent methods to study the diversity of viruses that infect *Acidobacteriota* in Arctic soils. Soil samples were obtained from meadows and fens in Kilpisjärvi, northern Finland. Using *Tunturibacter* (*Terriglobia*) strains originating from the same area (Kilpisjärvi) [46], we isolated five viruses, Tunturi 1-5, which to the best of our knowledge, represent the first reported isolates of viruses that infect *Acidobacteriota*. In addition to virus isolation, we explored virus-host interactions in Kilpisjärvi soils by bioinformatically predicting viral operational taxonomic units (vOTUs) from bulk metagenomes and linking them to their putative microbial hosts. A group of 125 vOTUs could be linked to *Acidobacteriota*, and among vOTUs that could be linked to putative hosts, this group was one of the most dominant ones across samples and especially in fens. *Terriglobia*-linked proviruses that were found among Kilpisjärvi vOTUs shared similarities with other proviruses predicted in *Acidobacteriota* strains that reside in various remote environments. Finally, Tunturi 1-5 showed genus- or higher-level links to Kilpisjärvi vOTUs, as well as vOTUs from other soils, but not with NCBI reference sequences. The viruses and vOTUs reported here provide a glimpse into the viral diversity associated with *Acidobacteriota* hosts in climate-critical Arctic soils.

## Materials and Methods

### Soil samples

The sampling sites are located in the oroarctic mountain tundra area in Kilpisjärvi, northwestern Finland (69.04°N, 20.79°E) [6,42]. The main vegetation cover in the sites were fens and meadows (Supplementary Table S1). Metagenomes from soil cores that were collected in July 2017 and July 2018 [6] were analyzed for viral sequences (see below, Identifying viral sequences in metagenomes). For virus isolation, fresh samples from the same area were collected in April 2021. Snow depth ranged from 72 to 99 cm and the mean air temperature was -2.9°C in Kilpisjärvi in April 2021 (https://en.ilmatieteenlaitos.fi/download-observations). Snow and frozen plant material were removed, and samples were chiseled from the top 5 cm of soil surface, spooned into ziplock bags and stored at 4°C. All sampling tools were sterilized with 70% ethanol prior usage.

### Bacterial strains and growth conditions

The 18 acidobacterial strains used as potential viral hosts in this study (Table S2) were previously isolated from Kilpisjärvi, Finland [44,46]. The tested strains belonged to four different genera of the class *Terriglobia*: *Tunturibacter*, *Granulicella*, *Acidicapsa*, and *Terriglobus*. The strains were grown in DSMZ medium 1284, containing 0.5 g L^−1^ glucose, 0.1 g L^−1^ yeast extract (Neogen, Lansing, USA), 0.1 g L^−1^ casamino acids (MP Biomedicals,Solon, USA), 0.04 g L^−1^ MgSO_4_ x 7 H_2_O, and 0.02 g L^−1^ CaCl_2_ x 2 H_2_O (https://www.dsmz.de/microorganisms/medium/pdf/DSMZ_Medium1284.pdf), pH 5.5. For plates and top-layer agar, 15 and 4 g of agar were added per 1 L, respectively. All culturing was done aerobically at room temperature (RT).

### Virus isolation

Infectious phage particles were extracted by resuspending soil samples in the DSMZ medium 1284 broth (approximately 1:3 ratio [wet weight]) and incubated with aeration at RT for 30 min. The supernatants (Table Eppendorf centrifuge, 2500 g, RT, 30 min) were filtered (0.22 μL LLG Syringe Filters Spheros filters) and the phage extract was applied to freshly grown host strains in the plaque assay: 100-150 μL of the supernatant were mixed with 300 μL of the host culture and 3 mL of the soft agar (46°C), and spread as a top layer on agar plates. The plates were incubated at RT and monitored for plaque formation regularly. The observed single plaques were picked up with a sterile pipette tip, resuspended in the DSMZ medium 1284 broth and subjected to the plaque assay, which was repeated three consecutive times to ensure the purity of virus isolates.

### Preparation of agar stocks

The top layers of semi-confluent plates were collected and mixed with the DSMZ medium 1284 broth (3 mL per plate), incubated aerobically at RT for 1 h, and the agar as well as cell debris were removed by centrifugation (F15-6×100y, 10000 g, 4°C, 30 min). The supernatant was collected, filtered (0.22 μL LLG Syringe Filters Spheros filters), and stored at 4°C. Stocks were titrated by the plaque assay method as described above.

### Virus host range testing

To determine viral host ranges, stocks were first subjected to the spot test. Plates having 300 μL of the bacterial liquid culture and 3 mL of the soft agar as a top layer were prepared, and 7-μL drops of undiluted and 100-fold diluted virus stocks were applied onto them. The drops of broth containing no virus sample were used as a negative control. The plates were incubated at RT and monitored for the growth inhibition. When inhibition was observed, the virus-host pair was additionally tested by the plaque assay with a range of dilutions to verify the spot test results.

### Virus purification

Viruses were precipitated from agar stocks by mixing with polyethylene glycol 8000 (PEG 8000, Thermo Scientific, final concentration 10% [w/v]) and NaCl (final concentration 0.5 M) and incubated with stirring at 4°C for 1 h. The pellets (F15-6×100y, 10000 g, 4°C, 30 min) were resuspended by adding the SM buffer (50 mM MES, pH 5.5; 100 mM NaCl; 8 mM MgSO4) in the amount of ∼1.5 % (v/v) of the original stock volume. If the resuspended pellets were highly viscous, the resuspension step was repeated with more SM buffer and in some cases, DNase (Stemcell Technologies) was also applied to reduce the viscosity (final concentration of 300 μg mL^−1^). The resuspended samples were further pelleted (F15-6×100y, 10000 g, 4°C, 10 min) and either the supernatant or both the supernatant and the pellet (separately) were used for purification by rate-zonal ultracentrifugation in 10-30 % (w/v) sucrose gradients in the SM buffer. The light-scattering zones observed after ultracentrifugation (Sorvall AH629 112142.4 g or TH641 103557.6 g, 10°C, 20-60 min) were collected and pelleted (Sorvall T1270, 113488.6 g, 4°C, 3 h). In case no clear light-scattering bands could be observed, the gradients were fractionated, and the fractions with highest virus titers were used for pelleting. Pellets were resuspended in 50-100 μL of the SM-buffer, titrated by plaque assay and stored at 4°C.

### Electron microscopy

The samples for transmission electron microscopy (TEM) were prepared by applying a drop of a PEG-precipitated or purified virus sample on the Mesh 200 cu grid for 1 min and rinsing it twice with ultrapure water. The samples were stained by applying a drop of Vitroease (Thermo Scientific) or 3% (w/v) uranyl acetate (pH 4.5) for 1 min, which was repeated twice. The images were taken with the JEOL 1400 electron microscope operating at 80 kV at the Electron Microscopy Unit, Institute of Biotechnology, University of Helsinki. The size of viral particles was measured with the ImageJ program [50]. The head size was measured as the distance between opposite vertices of icosahedral particles, except for the virus Tunturi 4, which had a prolonged icosahedral head. The number of particles used for head/tail measurements was 27/9, 10/7, 14/10, 27/12, and 5/11 for Tunturi 1-5, respectively.

### Genome sequencing and annotation of virus isolates

For the DNA extraction from virus stocks, the protocol by Santos [51] was used with modifications as described in [52]. The extracted DNA was purified using the GeneJET Genomic DNA purification Kit (Thermo Scientific). The purified DNA was sequenced using the Nextera XP kit and Illumina MiSeq (paired end, 325 bp + 285 bp) at the DNA Sequencing and Genomics Laboratory, Institute of Biotechnology, University of Helsinki. The quality of raw Illumina reads was assessed with FastQC v. 0.11.8 (https://www.bioinformatics.babraham.ac.uk/projects/fastqc/). Cutadapt v. 2.7 was used for removing adaptors and trimming reads (-q 30 -m 50) [53]. The virus isolate genomes were assembled using Spades v. 3.15.0 (-k 55,77,99,127) [54].

Geneious Prime v. 2021.2.2 (https://www.geneious.com) was used for the analyses of the viral genomes. Genome annotations were performed by Phold v. 0.2.0 [55] with Foldseek v. 9.427df8a [56], ProstT5 [57], and Colabfold v. 1.5 [58] as core dependencies, as well as PHROGs database [59]. In addition, DRAM-v v. 1.5.0 [60] was used for gene function predictions. Viral genome sequences produced circular maps, and in each virus isolate, ORFs were numbered starting from the ORF putatively encoding the terminase large subunit. The HHPred search against PDB_mmCIF70_8_Mar and SCOPe70_2.08 databases [61] was used to verify large teminase subunit predictions if contradictory predictions were produced by Phold and DRAM-v. tRNA genes were predicted using tRNAscan-SE v. 2.0 using bacterial search mode [62]. The programs fastANI v. 1.33 [63] and pyani v. 0.2.12 [64] were used for calculating average nucleotide identities (ANI) between the virus genomes. Overall nucleotide identities were calculated using Emboss Stretcher [65]. Intergenomic similarities were calculated with VIRIDIC [66]. Pairwise similarities between the genomes were visualized using Easyfig v. 2.2.2 with the BLASTn E-value threshold 0.001 [67].

The virus isolates genomes were searched against the IMG/VR v. 4 [68] using BLASTn v. 2.13.0 with the E-value threshold 1e-5. Similarities between genomes were visualized with Circoletto using the BLASTn E-value threshold 1e-5 [69]. To detect sequences related to virus isolates in Kilpisjärvi metagenomes, amino acid sequences from the five isolates were clustered with MMseqs2 v. 14 [70] to generate a non-redundant protein set (≥50% identity, ≥90% coverage). Quality-filtered metagenomic reads were then mapped to the set of non-redundant proteins with Diamond v. 2.1.6.160 [71] using the E-value threshold 1e-5.

### Identifying viral sequences in metagenomes

Previous metagenomic data from Kilpisjärvi fens and meadows soils [6] were analyzed for the presence of viral sequences. Raw reads were quality-checked and trimmed as described in [6]. Each of the 22 samples (Table S1) was assembled separately using metaSpades v. 3.14.1 (k-mers 55, 99, and 127) [54]. QUAST v. 5.0.2 [72] was used for the quality assessment of the assemblies. Quality-filtered metagenomic reads were mapped to the assemblies using Bowtie v. 2.4.1 [73]. For the identification of viral contigs, Virsorter v. 2.2 [74], PPR-Meta [75] and DeepVirFinder v. 1.0. [76] were used within the What-the-Phage pipeline [77]. These tools were recently benchmarked as having high sensitivity and precision [78]. The contigs identified as viral by the three tools with scores/p-values sum >0.75 were selected. In addition, geNomad v. 1.4.0 [79] was used for extracting viral sequences from the metagenomic assemblies. The predictions from the four tools were combined, their taxonomy was predicted using geNomad v. 1.4.0 [79], and the resulting contigs were checked for quality and completeness with CheckV v. 0.8.1 [80]. The contigs that were ≥ 5 kbp long or predicted as ≥ 50% complete (but not shorter than 1 kbp), had at least one viral gene, and no more than 1.5 host to viral genes ratio were selected for the final set of viral contigs. The set was dereplicated using CheckV -anicalc and -aniclust functions and all contigs within 95% average nucleotide identity and 85% alignment fraction were assigned to the same vOTU cluster, following the suggested standard thresholds [81]. For annotations and AMGs predictions in vOTUs, DRAM-v v. 1.5.0 [60] was used with the viral sequences preprocessed by the Virsorter v. 2.2.4 [82].

### Linking vOTUs to their putative hosts

Putative hosts for vOTUs were predicted with iPHoP v. 1.3.3 [83] using the minimum score cutoff of 90 and and 75 for the genus- and family-level predictions, respectively. Both the default database (iPHoP_db_Aug23_rw) and a custom database that included 796 metagenome-assembled genomes (MAGs) previously obtained from Kilpisjärvi metagenomes [6] were used. For proviruses, both proviral sequences and their corresponding larger contigs with remaining host regions were used as an input. For building the custom database, Kilpisijärvi MAGs were first classified using GTDB-tk v. 2.3.2 [84] with the GTDB release 214 (https://data.gtdb.ecogenomic.org/). One host prediction obtained for the vOTU o12215_NODE_6138, which clustered with Tunturi 3 in the VConTACT2 analysis (see below), was manually inspected and found to be based on a short entirely viral contig present in a MAG [85], so this prediction was discarded as unreliable.

Proviral vOTUs were predicted by both geNomad v. 1.4.0 [79] and CheckV v. 0.8.1 [80] and those assigned to *Acidobacteriota* hosts were further explored by comparing to previously reported acidobacterial proviruses [47] and UViGs that were retrieved from the IMG/VR v. 4 [68] with *Acidobacteriota* as a host. In addition, acidobacterial proviral vOTUs were searched against the NCBI nr database (discontinuous MegaBLAST, searches dated Feb-Mar 2024, E-value threshold 0.001, query coverage threshold 10%) [86], and for bacterial genomes returned as hits satisfying the thresholds, proviral regions were predicted with geNomad v. 1.4.0 [79] and further used for comparisons. The sequence similarities were analyzed using Circoletto with the BLASTn E-value threshold 1e-5 [69] and genome-to-genome comparisons were visualized with Easyfig v. 2.2.2 (BLASTn E-value threshold 0.001) [67]. High-quality vOTUs assigned to *Acidobacteriota* were annotated using Phold v. 0.2.0 [55] and DRAM-v v. 1.5.0 [60].

### vOTUs abundances

Quality-filtered metagenomic reads were mapped to acidobacterial vOTUs with Bowtie2 v. 2.5.3 [73] and the mapping output was sorted and indexed with SAMtools v. 1.16.1 [87]. CoverM v. 0.6.1 (https://github.com/wwood/CoverM) was then used to count the number of reads mapping to each vOTU, considering only matches with ≥95% identity and ≥75% coverage. CoverM v. 0.6.1 was also used to compute the fraction of each vOTU that was covered by at least one read (horizontal coverage, also known as breadth of coverage). Abundance values were normalized to reads mapped per kilobase of contig per million reads (RPKM), and the abundance of vOTUs with <50% horizontal coverage was set to zero. Statistical analyses were done with the package vegan v. 2.6-6.1 in R v. 4.4.2 (https://github.com/vegandevs/vegan, https://cran.r-project.org). Differences in vOTU abundances between meadows and fens samples were visualized using principal coordinates analysis (PCoA) and confirmed with permutational ANOVA (PERMANOVA) with 9999 permutations. For both, distances were computed using the binary (presence/absence) Jaccard dissimilarity metric. The contribution of soil physicochemical variables was also verified with PERMANOVA.

### Whole-genome comparisons using VConTACT2

Kilpisjärvi virus isolates genomes and vOTUs ≥10 kbp were used in the whole genome gene-sharing network analysis by VConTACT v. 2.0 [88] together with previously reported vOTUs from peat permafrost microbial communities from Stordalen Mire, Sweden [23], as well as acidobacterial proviruses identified in this study. The NCBI ProkaryoticViralRefSeq211-Merged database was used to resolve taxonomic clustering. The network was visualized with Cytoscape v. 3.9.1 [89].

## Results

### Virus isolation

Five new virus isolates, which we called Tunturi 1-5, were obtained from the Kilpisjärvi soil samples on *Tunturibacter psychrotolerans* and *T. empetritectus* strains originating from the same area (Table 1). Clear plaques of 1-5 mm in diameter were observed after 5-8 days of incubation. Stocks with titers reaching 10^8^-10^9^ plaque forming units per mL (PFUs mL^−1^) could be obtained for the isolates. The initial spot tests for the virus-host range with 18 acidobacterial strains previously isolated from Kilpisjärvi soils (Table S2) showed several inhibition zones, representing virus infections or bacterial growth inhibition by some chemical components of the virus stocks. Only one additional virus-host pair could be verified by plaque assay. The virus isolate Tunturi 4 could infect *Granulicella* sp. J1AC2, albeit with plating efficiency lower than the one with its original isolation host (10^7^ PFU mL^−1^ and 10^8^ PFU mL^−1^, respectively).

**Table 1.** Viruses isolated in this study.

| Virus name | Host strain | Sample site | Morpho-type*, size | Virus genome |  |  |  |  |
| --- | --- | --- | --- | --- | --- | --- | --- | --- |
|  |  |  |  | Length, bp | GC % | ORFs | tRNA genes | GenBank acc. number |
| Tunturi 1 | *Tunturibacter psychrotolerans* X5P2 | 12217 (fen) | Podo-, head 73.2 nm, tail 23.5 nm | 63,169 | 55.3 | 106 | 0 | PP887698 |
| Tunturi 2 | *Tunturibacter psychrotolerans* X5P2 | 12217 (fen) | Podo-, head 64.4 nm, tail 24.3 nm | 63,277 | 55.1 | 107 | 0 | PP885685 |
| Tunturi 3 | *Tunturibacter psychrotolerans* X5P2 | 181 (meadow) | Podo-, head 76.7 nm, tail 21.1 nm | 97,608 | 51.3 | 147 | 3 | PP885686 |
| Tunturi 4 | *Tunturibacter empetritectus* M8UP27 | 12222 (meadow) | Sipho-, head 94.1x54.2 nm, tail 204.1 nm | 88,042 | 55.4 | 115 | 1 | PP885687 |
| Tunturi 5 | *Tunturibacter psychrotolerans* X4BP1 | 12222 (meadow) | Myo-, head 112.7 nm, tail 167.0 nm | 308,711 | 58.4 | 350 | 43 | PP885688 |

\*Podo-, podovirus (short non-contractile tail); sipho-, siphovirus (long non-contractile tail); myo-, myovirus (long contractile tail). For viruses Tunturi 1-3 and 5, the head size was measured as the distance between opposite capsid vertices. The head of Tunturi 4 is a 94.1-nm long and 54.2-nm wide prolate icosahedron.

The electron micrographs of purified virus isolates showed that all five viruses displayed tailed particles with icosahedral heads, varying in size (Figure 1, Table 1). Viruses Tunturi 1-3 displayed icosahedral heads (∼64-77 nm) and short (∼21-24 nm) non-contractile tails of the podovirus morphotype. Tunturi 4 demonstrated a ∼94-nm long and a ∼54-nm wide elongated (prolate) icosahedral head and a ∼204-nm long flexible tail, featuring the siphovirus morphotype. Tunturi 5 had the largest head (∼113 nm) and a ∼167-nm long contractile tail typical for myoviruses, and both extended and contracted tail conformations were observed.

**Figure 1.**
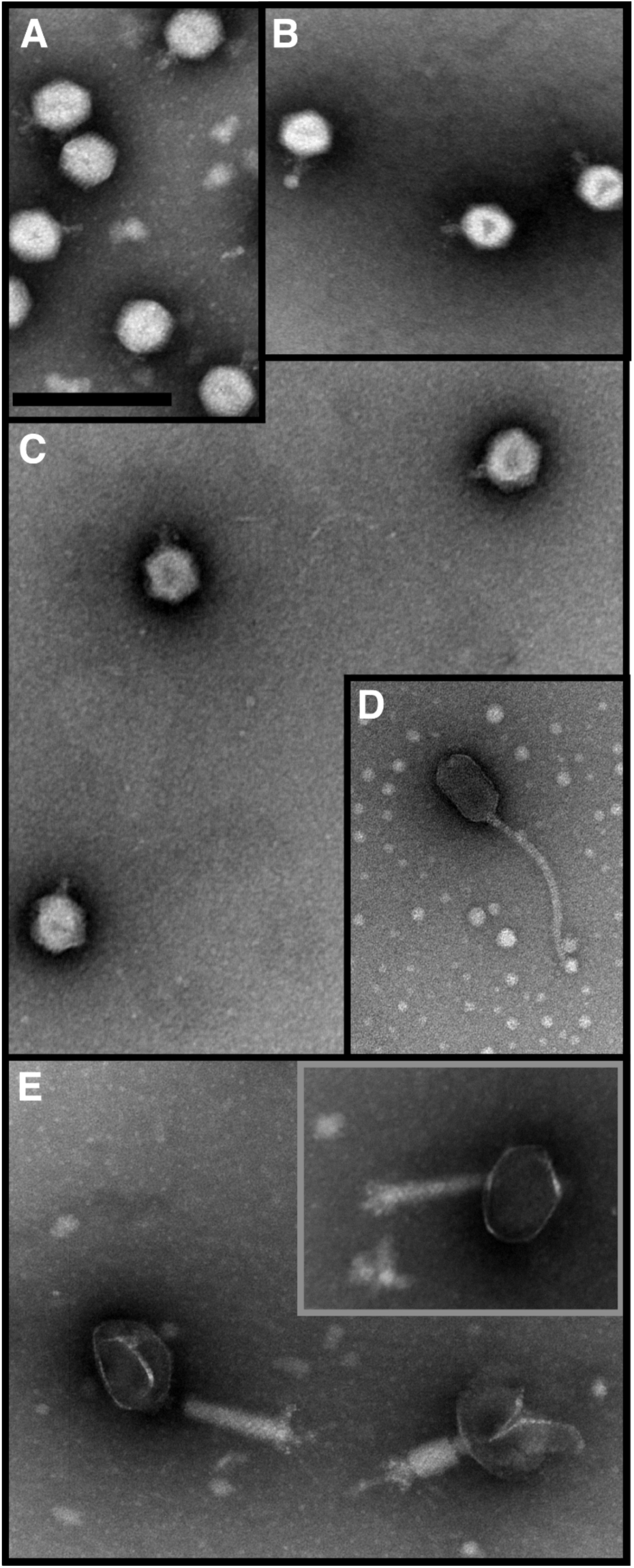
Electron micrographs of the five Tunturi virus isolates: (A) Tunturi 1, (B) Tunturi 2, (C) Tunturi 3, (D) Tunturi 4, (E) Tunturi 5. Virus particles were stained with Vitroease (in A, B, E) or 3% (w/v) uranyl acetate (in C and D). Scale bar in (A), 200 nm, for all sections.

### Genomic characterization of virus isolates

The genome length ranged from ∼63 to ∼98 kbp for the isolates Tunturi 1-4, while the Tunturi 5 genome was ∼309 kbp long (Table 1). The GC content varied from 51.3 to 58.4%, and the virus genomes were predicted to contain from 106 to 350 ORFs (Tables 1, S3), tightly packed in the genomes (1.1-1.7 ORFs/kbp, coding density 88-96%). The genomes of Tunturi 1 and Tunturi 2 were clearly related, having an ANI of 97.8% and overall nucleotide identity of 87.8% (Figure 2A). The other three genomes were not similar to one another. Based on the analyses of intergenomic similarities by VIRIDIC, all five isolates represent different species, but Tunturi 1 and Tunturi 2 clustered into the same genus.

**Figure 2.**
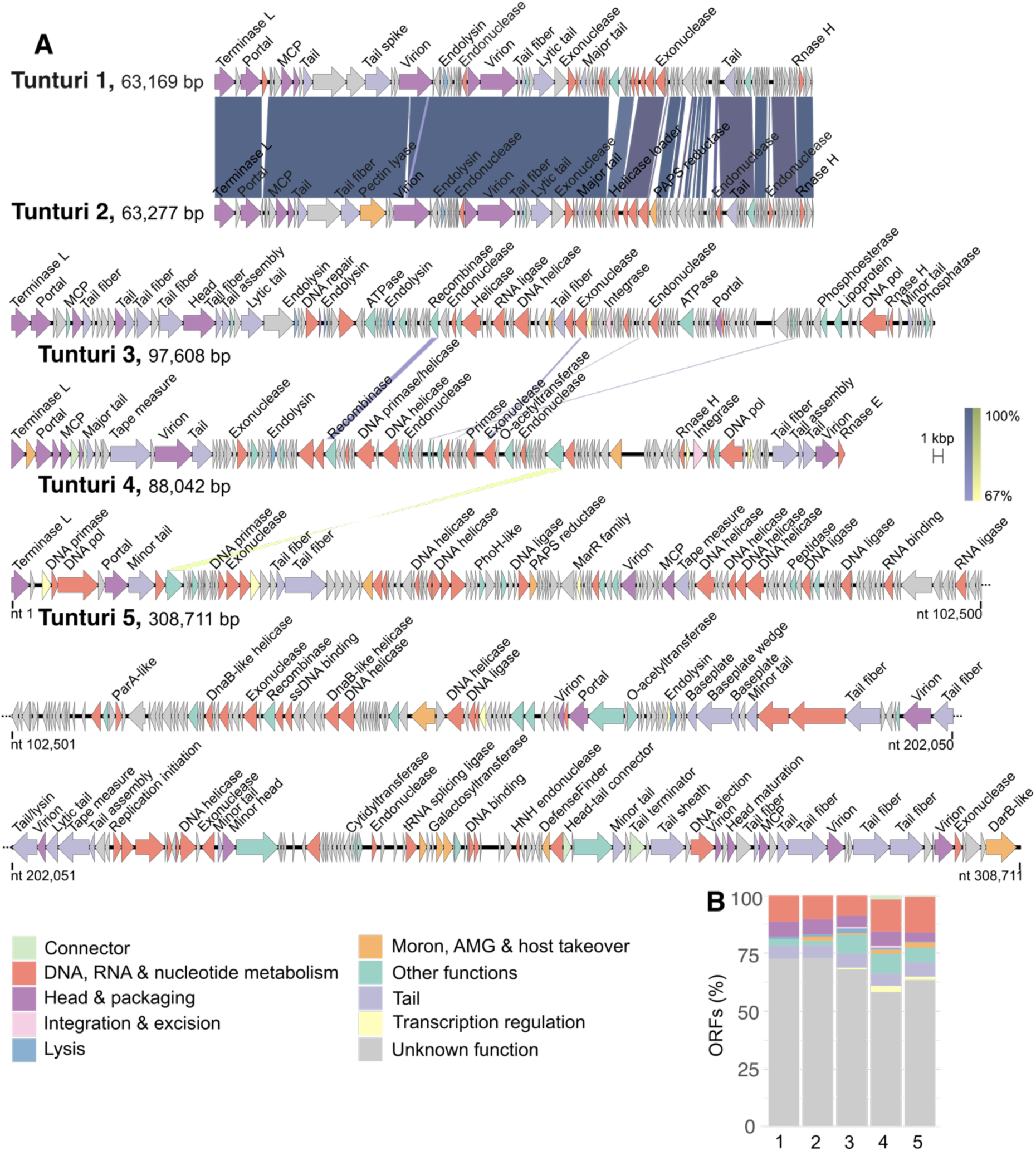
(A) Genomes of Tunturi 1-5. ORFs are shown as arrows and are colored according to the functional categories. Similarities between the genomes (BLASTn) are visualized by the shades of blue/yellow (direct/invert). (B) Distribution of ORFs according to the functional categories, 1-5 for Tunturi 1-5, respectively. The color code is the same for (A) and (B).

The majority of ORFs (58-73%) had no predicted functions (Figure 2B). The functions that could be predicted were related to DNA/RNA metabolism, virion structural elements, lysis or other functions typically found in phage genomes (Table S3). Lysis-related genes were predicted in all five isolates, while recombinase and/or integrase genes could be annotated in Tunturi 3-5 genomes. In addition, ORF functions categorised by Phold as “Moron, AMG and host takeover” were predicted for Tunturi 2-5. In Tunturi 5, eight such ORFs were found: glycosyltransferase (gp35), phosphoadenosine phosphosulfate reductase (gp62), ADP-ribosyltransferase exoenzyme toxin (gp205), ribosomal protein S6 glutaminyl transferase (gp297), galactosyl transferases (gp300 and gp301), DefenseFinder protein (gp317), and DarB-like antirestriction (gp350). In the HHPred search against PDB_mmCIF70_8_Mar and SCOPe70_2.08 databases, Tunturi 5 DefenseFinder protein gp317 got two best hits to BrxU, GmrSD-family Type IV restriction enzyme (acc. no. 7P9K, 99.8% probability) and SspE protein (6JIV, 99.6%), i.e., enzymes involved in bacterial systems of protection from viral infections. A few other >95% probability HHPred hits included enzymes involved in chromosome segregation in bacteria.

Tunturi 1 and 2 genomes had no predicted tRNA genes. Tunturi 3 encoded three tRNAs (Asn, Phe and unknown) and Tunturi 4 encoded one tRNA (unknown). In contrast, Tunturi 5 genome contained 43 tRNA genes having the anticodon sequences of 20 different amino acids: Leu (4 tRNA genes), Cys (1), Tyr (1), Ser (3), Asn (1), Gln (2), Gly (2), Thr (3), Pro (3), His (1), Ala (3), Phe (1), Arg (5), Trp (1), Asp (1), Met (1), Ile (2), Lys (2), Glu (2), Val (1) and three unknown ones. In addition, Tunturi 5 was predicted to encode proteins involved in tRNA processing: tRNA nucleotidyltransferase (gp279) and a tRNA splicing ligase (gp296).

When Tunturi genomes were searched against the IMG/VR database, many hits to soil metagenomes could be retrieved. Longer stretches of similarities (typically ≤ 70% nt identity) were observed, for example, against Arctic soil microbial communities from a glacier forefield, Greenland (Tunturi 1 and 2) and peat permafrost microbial communities from Stordalen Mire, Sweden (Tunturi 3 and 5) (Figure S1, Table S4). However, the hits were not limited to the Arctic and included tropical soils from Puerto Rico (Tunturi 1 and 2) and soils from Indiana, Colorado and Washington in the USA (Tunturi 3 and 4). When comparing Tunturi viruses against Kilpisjärvi metagenomes, a small number of reads (up to 0.025%) could be mapped to Tunturi protein coding sequences (CDSs, up to 50% CDSs per viral genome) (Figure S2). Tunturi genomes were further compared to Kilpisjärvi vOTUs in a whole-genome analyses using VConTACT2 (see below).

Tunturi genomes were further tested with iPHoP to assess the performance of the tool on viruses with a known host. For Tunturi 1, 2, and 5, hosts were predicted from the family *Acidobacteriaceae* (class *Terriglobia*). Tunturi 1 and 2 could be further predicted with a host from the genus *Edaphobacter.* These matches were meaningful, taking into account that the isolation hosts of these viruses, i.e., *T. psychrotolerans* strains, were indeed formerly associated with the genus *Edaphohacter.* The new nomenclature has been proposed only recently and thus has not yet been reflected in the Genome Taxonomy Database (used for MAGs classification when including them into the custom iPHoP database) and the default iPHoP database. No hosts were predicted for Tunturi 3 and 4 using iPHoP. Having host predictions for three viruses out of five in this test was consistent with the expected tool performance on soil viruses [83].

### Metagenome-derived vOTUs

The 22 assembled metagenomes from fen and meadow Kilpisjärvi soil samples produced 491,604-9,096,281 contigs per sample (Table S5). The reads were mapped back to the metagenomic assemblies with a 31-79 % overall alignment rate (Table S5). From these metagenomes, 1881 vOTUs were predicted, from which 184 were of medium or high quality, including 46 vOTUs predicted as ≥90% complete virus genomes (Table S6). The vOTUs ranged from 2.3 to 208 kb in length (median 8.75 kb, note that each vOTU consisted of a single contig), and 794 vOTUs were ≥10 kbp. Based on the geNomad taxonomic assignments, the majority of vOTUs (1843 = 98%) were classified as dsDNA tailed viruses belonging to the class *Caudoviricetes*, from which four vOTUs could be further classified to the family *Herelleviridae* and one to *Straboviridae*. Among other predicted classes, *Tectiliviricetes* (3), *Malgrandaviricetes* (2), *Polintoviricetes* (1), *Megaviricetes* (1), *Faserviricetes* (1), and *Herviviricetes* (1) were found. In addition,1 vOTU was assigned to the kingdom *Bamfordvirae* with no further levels of classification and 28 vOTUs stayed unclassified. vOTUs classified as *Caudoviricetes* dominated across all samples (Figure S3).

### Virus-host linkages

In total, 722 vOTUs could be linked to putative hosts using iPHoP (Figure 3A, Table S6). From all matches, 418 were found from both default and custom databases, 166 only from the default database, and 138 only from the custom one. Thus, adding Kilpisjärvi MAGs to the iPHoP database noticeably increased the number of predictions, highlighting potential connections to local microbial hosts. For 687 vOTUs, the host could be predicted at least at the phylum level. The largest predicted group of hosts was the phylum *Pseudomonadota* (162), followed by *Actinomycetota* (159) and *Acidobacteriota* (125). In addition to bacteria, 33 archaeal hosts were also predicted, most of which belonged to the phylum *Halobacteriota* (17).

**Figure 3.**
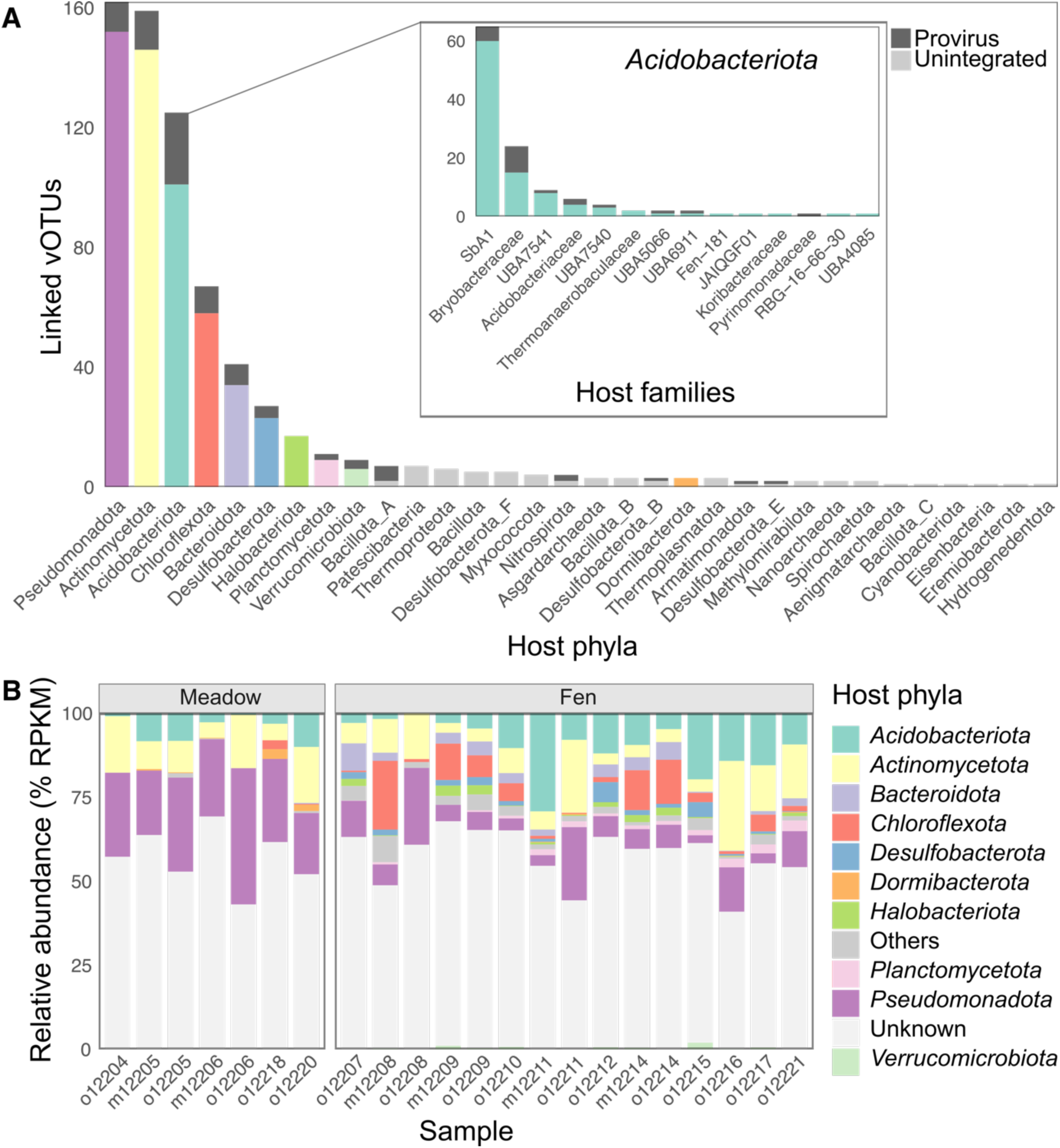
Host predictions for vOTUs obtained in this study. (A) Total number of vOTUs linked to different phyla and. The inset shows the distribution of predicted *Acidobacteriota* hosts at the family level. Proviruses stacks are highlighted with dark gray. (B) Relative abundance of vOTUs assigned to different host phyla across samples (10 most abundant ones are shown colored). Color codes are the same for (A) and (B).

Based on relative abundance (Figure 3B), most of the samples were dominated by vOTUs with unknown hosts. However, vOTUs assigned to *Pseudomonadota* and *Actinomycetota* were abundant in many samples in both meadows and fens. *Pseudomonadota*-linked vOTUs were especially dominant in meadows. *Acidobacteriota* vOTUs were present in almost all samples, but their highest relative abundances were in fens. *Chloroflexota* vOTUs also constituted a large group in fens. Other vOTUs groups were noticeably less abundant. According to PCoA, *Acidobacteriota*-assigned vOTUs, as well as the whole set of vOTUs identified in this study, formed different communities between Kilpisjärvi fens and meadows (Figure S4). PERMANOVA analysis showed that environmental variables that contributed to these differences most were soil moisture, SOM, C, N content and C:N ratio, but not soil layer (organic/mineral) or pH.

### High-quality vOTUs linked to *Acidobacteriota*

Among acidobacterial hosts, the families *SbA1* (65) and *Bryobacteraceae* (24) were represented the most (Figure 3A inset). Five *Acidobacteriota* vOTUs were of high quality: 96-100% complete, 44-59 kbp, all classified as *Caudoviricetes*, and putatively assigned to the host families *SbA1* and *Acidobacteriaceae*. In these five vOTUs, most of the predicted ORF functions were those involved in head and packaging, tail structures, and DNA/RNA metabolism (Figure S5, Table S7). Three out of five vOTUs had putative lysis genes and three had integrase genes. In each vOTU, moron, AMG and host takeover proteins could be predicted, including galactosyl and glycosyl transferases, phosphoadenosine phosphosulfate reductase, DarB-like antirestriction, membrane protein, polysaccharide deacetylase, ferredoxin, acyl carrier protein, Ren-like exclusion protein and GtrB-like O-antigen conversion protein. Many ORFs (47-66%) remained with unknown functions (Figure S5, Table S7).

### Predicted *Acidobacteriota* proviruses

In total, 114 vOTUs were identified as proviruses, and for 90 of them, bacterial hosts could be predicted, including eight predictions only at the domain level (Table S6). No archaeal hosts were predicted for proviral vOTUs. The three largest groups among predicted proviral hosts were *Acidobacteriota* (24 vOTUs assigned), *Actinomycetota* (13), and *Pseudomonadota* (10) (Figure 3A). In acidobacterial proviruses predicted in this study, two larger groups with shared sequence similarity and gene order could be identified: group (i) (Figure 4) and group (ii) (Figure S6). All Kilpisjärvi vOTUs belonging to these two groups were classified as *Caudoviricetes*.

**Figure 4.**
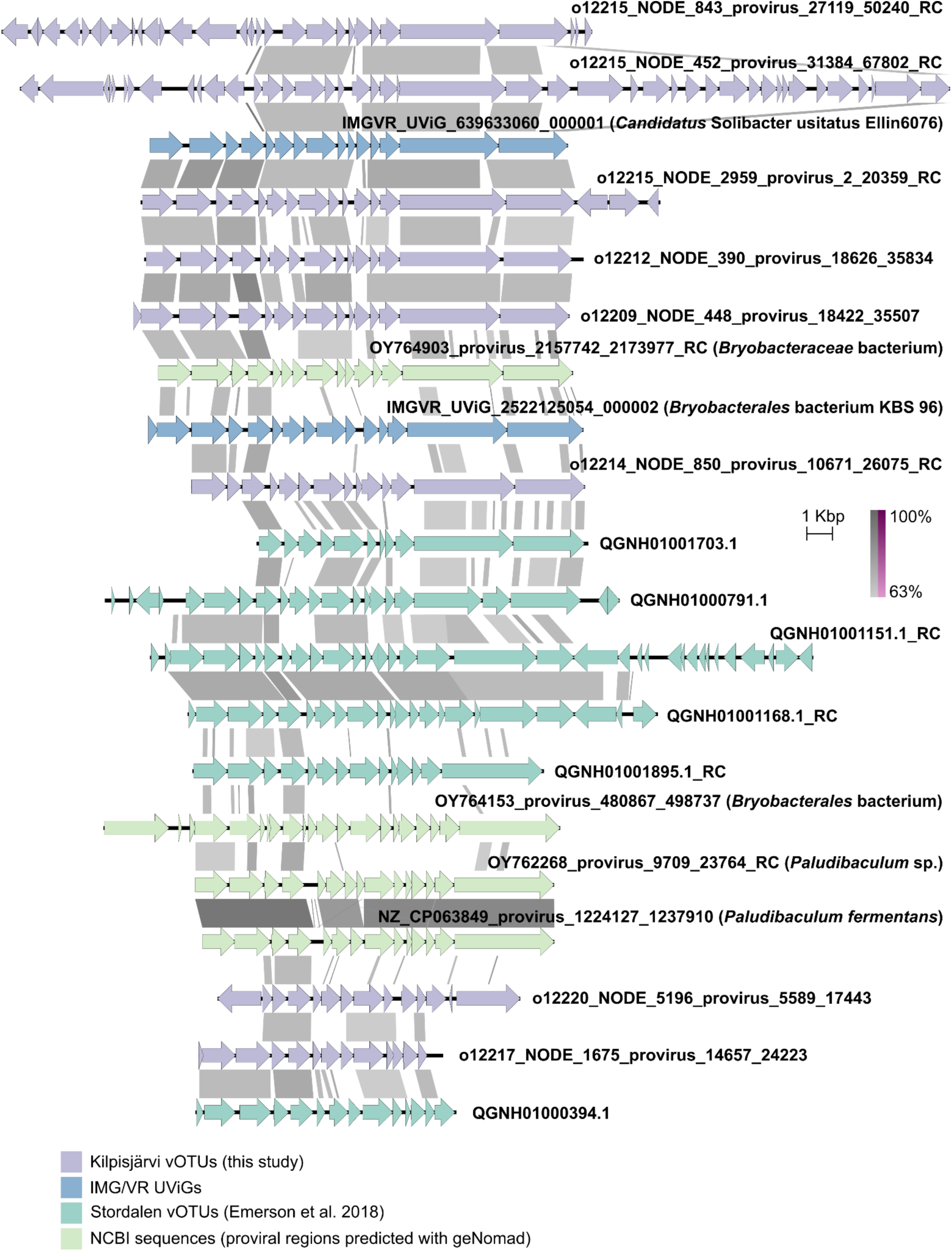
Putative *Acidobacteriota*-associated proviruses identified in this study and related IMG/VR UViGs, vOTUs reported in [23], and NCBI references. ORFs are shown as arrows and similarities between genomes (BLASTn) are in shades of gray (direct) or purple (invert). RC, reverse complement.

In addition to Kilpisjärvi *Acidobacteriota*-linked proviral vOTUs, group (i) (Figure 4) included Stordalen vOTUs [23] from the same VConTACT2 cluster (see below), and with iPHoP, all these Stordalen vOTUs could be putatively assigned to acidobacterial hosts from the family *Bryobacteraceae.* Two UViGs from the IMG/VR database also belonged to the group: IMGVR_UViG_639633060_000001 (*Candidatus* Solibacter usitatus Ellin6076) and IMGVR_UViG_2522125054_000002 (*Bryobacterales bacterium* KBS 96). In addition, a few proviral regions predicted in acidobacterial genomes retrieved from NCBI shared similarities with these proviruses. These NCBI references included *Paludibaculum fermentans* P105 (NZ_CP063849) isolated from a littoral wetland of a boreal lake on Valaam Island (Karelia, Russia) [90] and three MAGs annotated as *Bryobacteraceae* bacterium (OY764903), *Bryobacterales* bacterium (OY764153), and *Paludibaculum* sp. (OY762268) originating from freshwater ciliate *Cyclidium porcatum*, UK (genome assembly GCA_963668605.1), marine sponge *Geodia parva*, Norway (GCA_963667885.1), and freshwater ciliate *Heterometopus palaeformis*, Croatia (GCF_963665245.1) metagenomes, respectively.

Group (ii) (Figure S6) included Kilpisjärvi *Acidobacteriota*-assigned proviral vOTUs and a few (6-18% complete) acidobacterial vOTUs from the same dataset that were not recognised as proviruses by geNomad or CheckV. These vOTUs, however, formed one cluster in the VConTACT2 analysis. In addition, the group also included Stordalen vOTUs [23] from the same cluster. All these Stordalen vOTUs could be predicted with acidobacterial hosts, including families *SbA1*, *Koribacteraceae*, and *Acidobacteriaceae*, i.e., all belonging to the order *Terriglobales*. The group members shared similarities with the previously reported *Acidobacteriota* proviruses, like *Candidatus* Koribacter versatilis Ellin 345 [47]. Similar proviral regions were also predicted in *Granulicella* sp. WH15 (NZ_CP042596) isolated from decaying wood in association with the white-rot fungus *Hypholoma fasciculare* (Netherlands) [91] and a MAG annotated as *Granulicella* sp.(OY843766) from a lichen *Cladonia squamosa* metagenome, UK (genome assembly GCA_947623385.2, [92]).

### AMGs predicted in vOTUs

Using DRAM-v, 65 AMGs could be predicted in 58 Kilpisjärvi vOTUs (Table S8). Most of the detected hits were one per vOTU, but five vOTUs had more than one AMG predicted. All of the vOTUs, for which AMGs were predicted, were assigned to the class *Caudoviricetes* and 25 vOTUs had putative hosts, including two archaeal ones. Overall, the predicted AMGs categories included transporters, carbohydrate utilization, organic nitrogen transformation, and miscellaneous functions. From 21 putative AMGs involved in carbon utilization, most (19) were glycosyl transferases (GT2), but also two glycoside hydrolases involved in xyloglucan oligo cleavage were predicted. Six vOTUs bearing carbon utilization AMGs could be linked to the hosts from a few different phyla: *Chloroflexota* (families *EnvOPS12*, *Fen-1039*), *Halobacteriota* (*Methanosarcinaceae*), *Pseudomonadota* (*Gallionellaceae*, *Nitrosomonadaceae*), and *Actinomycetota* (*Mycobacteriaceae*). One vOTU with a predicted AMG (thymidylate synthase involved in pyrimidine deoxyribonuleotide biosynthesis) could be linked to the *Acidobacteriota* (*SbA1*) host.

### Whole genome comparisons

In the whole genome gene-sharing network analysis by VConTACT2 with NCBI ProkaryoticViralRefSeq211-Merged database (Figure 5, Table S9), Tunturi 1 and Tunturi 2 clustered together. Tunturi 5 was a singleton. Tunturi 3 clustered with one vOTU from the Kilpisjärvi dataset (o12215_NODE_6138, predicted as *Caudoviricetes*) and three vOTUs from the Stordalen Mire dataset (QGNH01000767.1, QGNH01001143.1, QGNH01001870.1) [23] (Figure S7). Tunturi 4 also clustered with one Stordalen vOTU (QGNH01000831.1) [23]. Using iPHoP, no hosts could be reliably predicted for o12215_NODE_6138 (see Materials and Methods) and these four Stordalen vOTUs.

**Figure 5.**
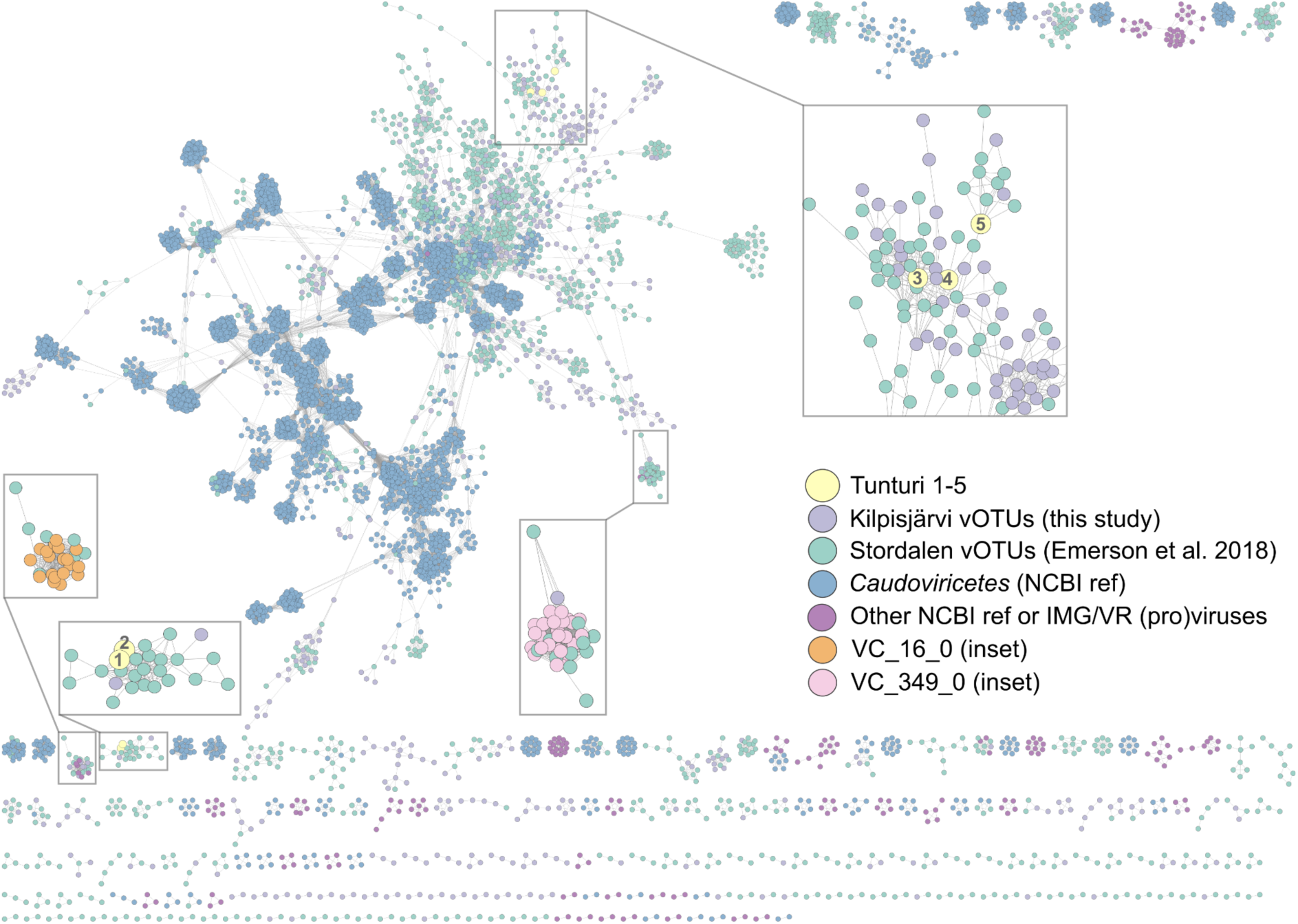
VConTACT2 gene-sharing network showing virus sequences as dots and connections between genomes as lines. The sequences are color-coded (see the color key). The insets show Tunturi viruses, highlighted with numbers 1-5 for Tunturi 1-5, respectively; and virus clusters VC_16_0 and VC_349_0 comprising putative proviruses shown in Figures 4 and S6, respectively.

From 794 Kilpisjärvi vOTUs used in the VConTACT2 analysis, 375 got clustered, while the others were singletons, outliers or couldn’t be confidently placed into a single cluster (overlapped). From the clustered Kilpisjärvi vOTUs, 186 vOTUs shared clusters only with Stordalen vOTUs and 181 vOTUs clustered only with other Kilpisjärvi vOTUs. Four clusters were shared by Kilpisjärvi and Stordalen vOTUs and NCBI sequences. Four Kilpisjärvi vOTUs clustered with NCBI reference sequences only.

From 58 Kilpisjärvi *Acidobacteriota*-linked vOTUs that were included into the analysis, 32 could be clustered. Almost all (31) these vOTUs shared clusters with Stordalen vOTUs. From the five high-quality *Acidobacteriota* vOTUs, o12205_NODE_77 was an outlier; o12215_NODE_1195 and o12215_NODE_1196 clustered together and with other Kilpisjärvi and Stordalen vOTUs; m12209_NODE_338 clustered with Stordalen vOTUs only; and m12211_NODE_265 with another Kilpisjärvi vOTU only. All Stordalen vOTUs that clustered with Kilpisjärvi high-quality *Acidobacteriota* vOTUs could also be predicted with *Terriglobales* hosts using iPHoP. Among *Acidobacteriota* proviruses included into the VConTACT2 analysis, one was outlier, two were singletons, and the rest belonged to four clusters, which all contained Stordalen vOTUs linked to *Terriglobia* with iPHoP. Two larger clusters, VC_16_0 and VC_349_0, included proviruses from groups (i) and (ii), respectively, which were described above.

## Discussion

In this study, we presented five viral isolates and 1881 metagenome-assembled vOTUs from Arctic tundra soils, including 125 vOTUs bioinformatically linked to *Acidobacteriota* hosts. To the best of our knowledge, the viruses isolated on *Tunturibacter* strains here are the first *Acidobacteriota*-infecting virus isolates reported. Following the suggested criteria for genome-based phage taxonomy [93], we propose that the five isolates described in this study represent five different species, belonging to the class *Caudoviricetes*. Based on the set of strains tested in this study, the Tunturi viruses have a narrow host range, which may be limited to their original isolation hosts, *Tunturibacter* strains, or may include other genera, as in case of Tunturi 4 being able to infect *Granulicella* sp. J1AC2. These virus-host pairs could be used as laboratory models for future studies, including developing genetic tools for the research on *Acidobacteriota*, which play important roles in key ecological processes in soil and other ecosystems.

From the five isolates, Tunturi 5 had the largest head and the largest genome, ∼309 kbp. Myoviruses tend to have larger genomes than other tailed phages, but the genome length of >200 kbp qualifies Tunturi 5 as a jumbo phage [94]. Jumbo phages have been isolated from various environments, but more frequently from water environments, rather than soils [94]. However, metagenomics-based studies reveal the presence of jumbo phages across ecosystems [95]. It has been observed that larger phage genomes tend to have more tRNA genes [95,96]. Indeed, Tunturi 5 had 43 tRNA genes and also encoded enzymes putatively involved in tRNA modification and maturation. Larger sets of tRNA genes in larger phage genomes seem to represent codons that are highly used by phages, while being rare in host genomes, and thus, may contribute to higher efficiency in phage protein translation [96]. In addition, the large genome of Tunturi 5 contains several putative moron, AMG and host takeover genes, which may specifically contribute to the mechanisms of virus-host interactions but need experimental validation. Having laboratory isolates available makes it possible to link sequences with processes and genes with functions. For example, single-step life cycle experiments could be developed, the patterns of gene expression analysed, knock-out mutant generated to determine essential genes, and gene functions confirmed with proteomics. Resolving molecular details of the interactions between *Acidobacteriota* and their viruses could help understand factors affecting their dynamics and ecosystem functions, in particular, in climate-critical Arctic tundra soils.

The five Tunturi isolates displayed tailed particles, with all three different tail types, and the vast majority of metagenomic Kilpisjärvi vOTUs were classified as tailed viruses belonging to the class *Caudoviricetes*. Microscopy-based studies have shown that different soil types may be dominated by different virus morphotypes [97,98]. In omics studies, *Caudoviricetes* typically represent a large fraction of those dsDNA viral populations that can be classified [23,25,68,99,100]. It remains to be seen if the vOTUs identified here represent active members of viral communities in Kilpisjärvi soils. In Alaskan peat soils studied under simulated winter conditions with stable isotope probing targeted metagenomics, active viral populations constituted a large portion of the whole viral communities [49]. In Stordalen Mire soils, 58% of vOTUs predicted from metagenomes were detected also in metatranscriptomes, thus being presumably active [23].

About 62% of Kilpisjärvi vOTUs stayed with unknown hosts, which is in line with the iPHoP benchmark, where 50-70% of virus genomes are expected to have no host prediction in soil [83]. From the predicted hosts, most numerous predictions were for the phyla *Pseudomonadota*, *Actinomycetota*, and *Acidobacteriota*, i.e., bacteria that are abundant and active in Arctic soils [3,6,42,101,102], as well as in other ecosystems [39]. *Acidobacteriota* were one of the largest groups of hosts predicted for viral populations in Kilpisjärvi, similarly to other (sub)arctic soils: Stordalen Mire soils [23,48] and the active layer of Alaskan tundra soils at CiPEHR warming experiment [25]. vOTUs linked to *Acidobacteriota* were abundant across samples but formed different communities in fens and meadows. Soil moisture, SOM, C, N content and C:N ratio contributed to the differences in viral communities across the two types of sites, similarly to the factors driving *Acidobacteriota* in Kilpisjärvi [6]. We could not detect the effect of pH on viral communities in the tundra samples studied here, all of which were rather acidic (pH 4.7-6.5). It has been shown that, for example, soil moisture [97,103,103,104], C and N content [25], soil depth [100], and pH [23] can be factors in modulating viral abundances and life styles. However, it is difficult to assess the causal relationships between environmental variables and dynamics in viral and host populations, as environmental parameters can be interconnected [105] and changes in viral communities can be linked to the processes going on with their hosts rather than directly environmental impacts. Our current understanding of soil ecology still lacks a clear view of how multiple biotic and abiotic factors collectively drive viral communities in soil [16,22].

The Tunturi viruses demonstrated lytic infection cycles, but their genomes contained ORFs for putative integrases and recombinases. Near-complete Kilpisjärvi vOTUs linked to *Acidobacteriota* also contained recombinase/integrase-encoding ORFs and ∼20% of Kilpisjärvi *Acidobacteriota*-linked vOTUs were recognised as proviruses. Lysogeny is common in soils [106,107], and the majority of prophages predicted in acidobacterial genomes have been found for strains originating from soil [47]. Based on the known/predicted hosts, two groups of *Acidobacteriota*-linked proviruses identified in this study seem to be specific for *Bryobacterales* (group (i)) and *Terriglobales* (group (ii)). The latter one is related to previously detected proviruses, while the former one is newly reported in this study. A large set of *Terriglobia*-associated putative proviruses described here includes vOTUs from northern soils, as well as proviral sequences in acidobacterial strains/MAGs originating from distant geographical locations and various environments, such as soil and ciliate-, sponge-, and lichen-associated biomes. These proviruses seem to be fairly widespread in *Acidobacteriota* across various environments and their diversity is yet to be uncovered.

Only a very small fraction of Kilpisjärvi metagenomic reads could be mapped to Tunturi CDSs, which is not unexpected taking into account that bulk metagenomes typically contain only a small number of viral sequences compared to cellular ones [108] and cultivable viruses may be in fact rare in natural communities [109]. Nonetheless, multiple genus- or higher-level links between Tunturi viruses, Kilpisjärvi vOTUs and Stordalen vOTUs could be found with the network-based whole genome gene-sharing profiles by vConTACT2. Using IMG/VR, viral sequences related to the Tunturi viruses could be also detected in metagenomes from other Arctic and temperate soils, suggesting some shared viral diversity and functions across soils. Global species-level sequence conservation across soil habitats has been observed when viromes from boreal peatland in northern Minnesota were compared with the PIGEON database having viral sequences from diverse ecosystems [100]. Similarly, shared viral clusters have been reported when comparing viromes from Alaskan permafrost and Stordalen Mire [25]. It is, however, unclear whether the observed patterns truly represent biological diversity or are biased because of the available (and yet limited mostly to peats) deeply sequenced virome data [100]. Largely unknown viral sequences detected in soils highlight a need for more extensive sampling to better understand viral functions and contributions to ecosystem-wide nutrient cycling processes, especially in the climate-wise vulnerable Arctic region.

## Ethics approval and consent to participate

Not applicable.

## Consent for publication

Not applicable.

## Availability of data and materials

Tunturi genome sequences are available from the NCBI with the following accession numbers: PP887698, PP885685-PP885688 for Tunturi 1-5, respectively. Kilpisjärvi vOTUs sequences described here can be downloaded from https://figshare.com/articles/dataset/Kilpisj_rvi_vOTUs/25976386?file=46841947.

## Authors’ contributions

TD, ISP, MKM, BED, SR and JH contributed to the study conception and design. HM, ISP and JH collected soil samples. TD and HM performed laboratory experiments and collected the data. TD and ISP analysed the data and designed figures. BED, SR and JH provided guidance on data visualization and interpretation. TD drafted the manuscript and all authors reviewed it.

## Acknowledgements

We acknowledge Riina Ihonen for technical assistance. We thank Prof. Max Häggblom for valuable discussions about *Tunturibacter* classification. We acknowledge Electron Microscopy Unit (EMBI), DNA Genomics and Sequencing core facility, and Biocomplex unit (member of Instruct-ERIC Centre Finland, FINStruct, and Biocenter Finland), Helsinki Institute of Life Science (HiLIFE), University of Helsinki. We also acknowledge CSC – IT Center for Science, Finland, for computational resources as well as for technical support.

## Supplementary Figures

**Figure S1.**
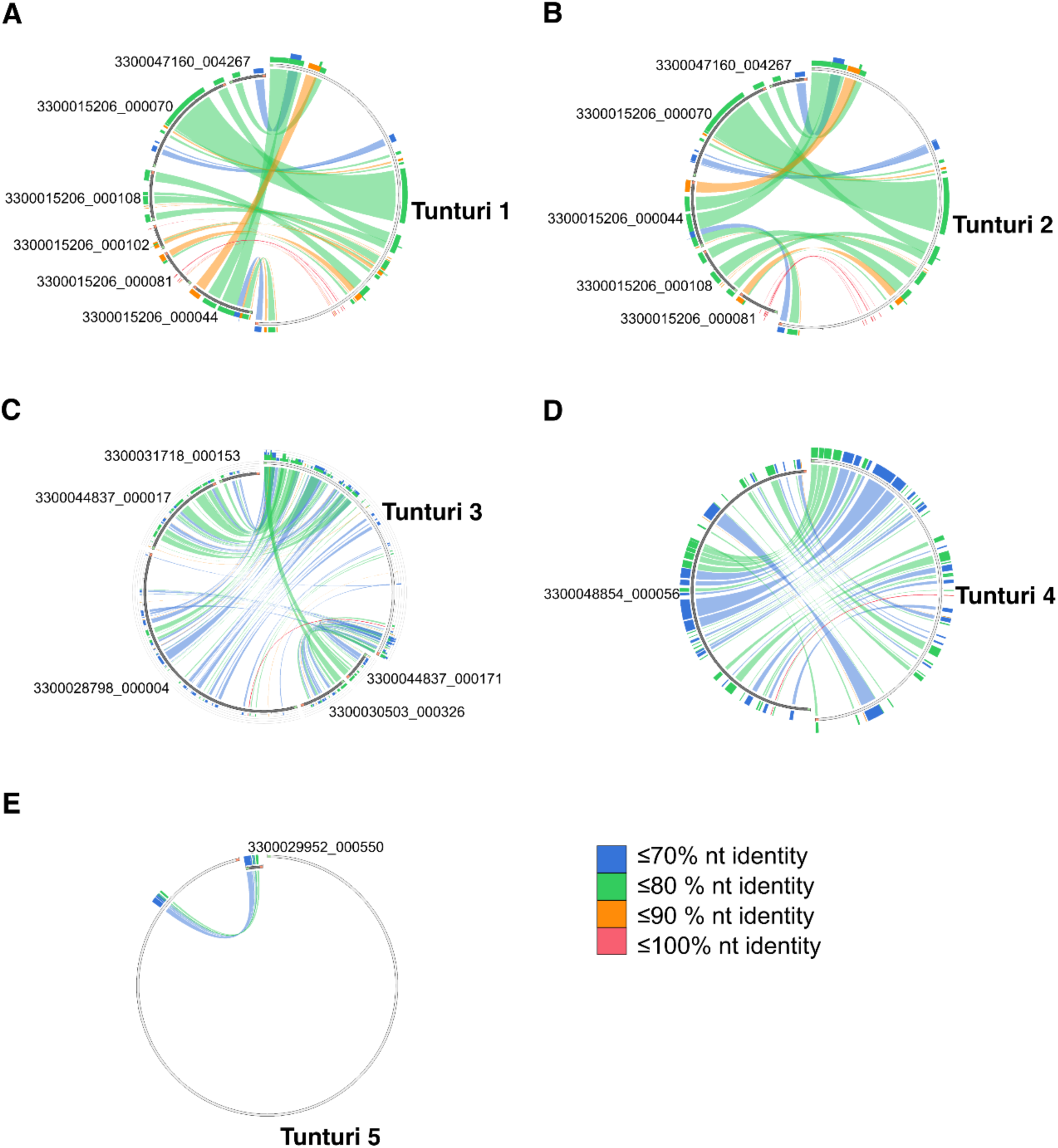
Similarities between the genomes of Tunturi 1-5 viruses and UViGs retrieved from the IMG/VR database. UViGs IDs are labeled skipping the “IMGVR_UViG_” prefix, for more information about UViGs, see Table S4. Ribbons are colored by % identity (see the key). Minimum and maximum % identities: (A) 64.71 and 100.00, (B) 64.96 and 100.00, (C) 65.11 and 95.83, (D) 64.79 and 90.32, (E) 64.69 and 79.45. Orientation is clockwise for all sequences.

**Figure S2.**
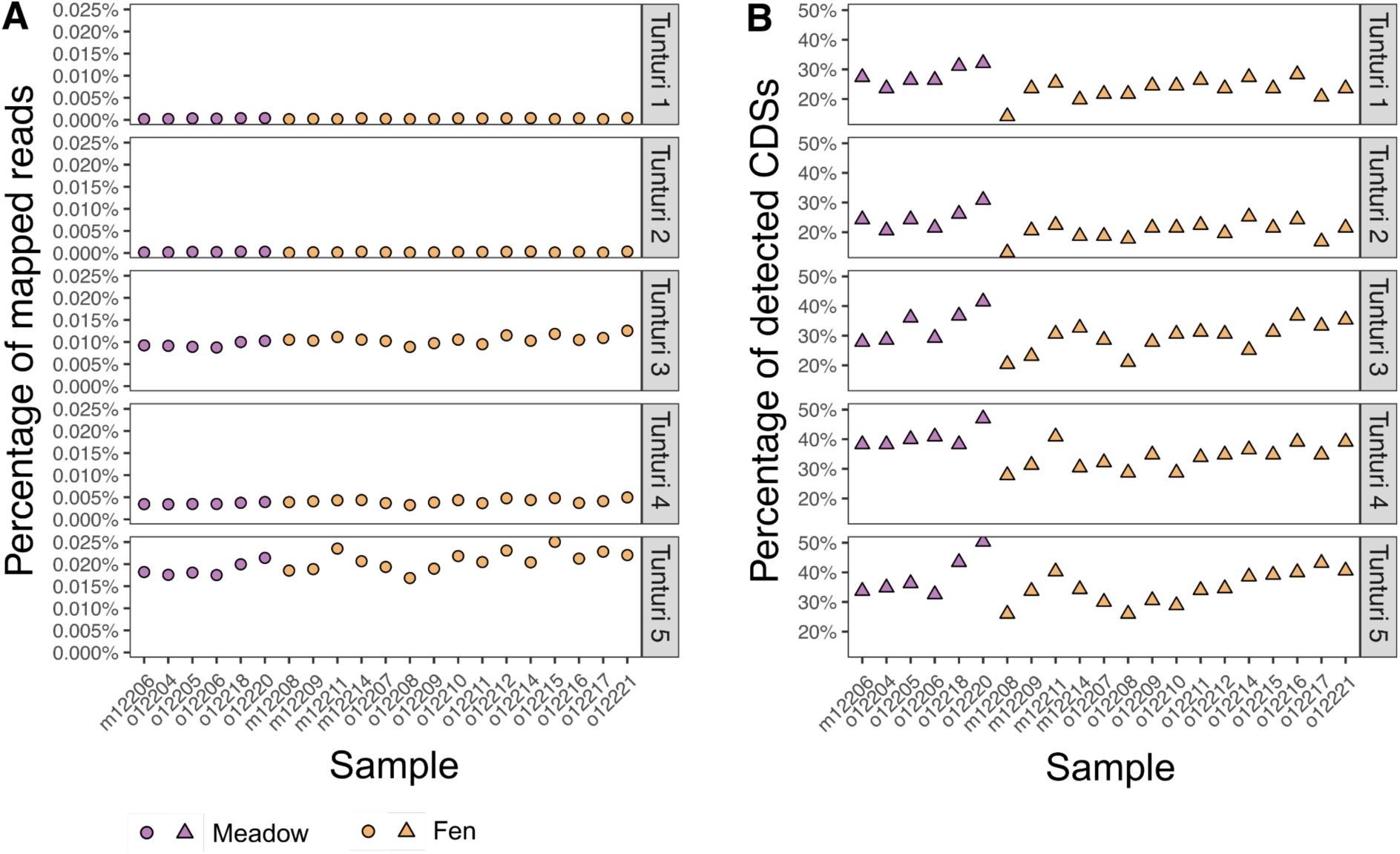
Distribution of the Tunturi 1-5 isolates across the 22 Kilpisjärvi meadow and fen metagenomes shown as (A) the percentage of reads mapped to their CDSs and (B) the percentage of detected CDSs per virus.

**Figure S3.**
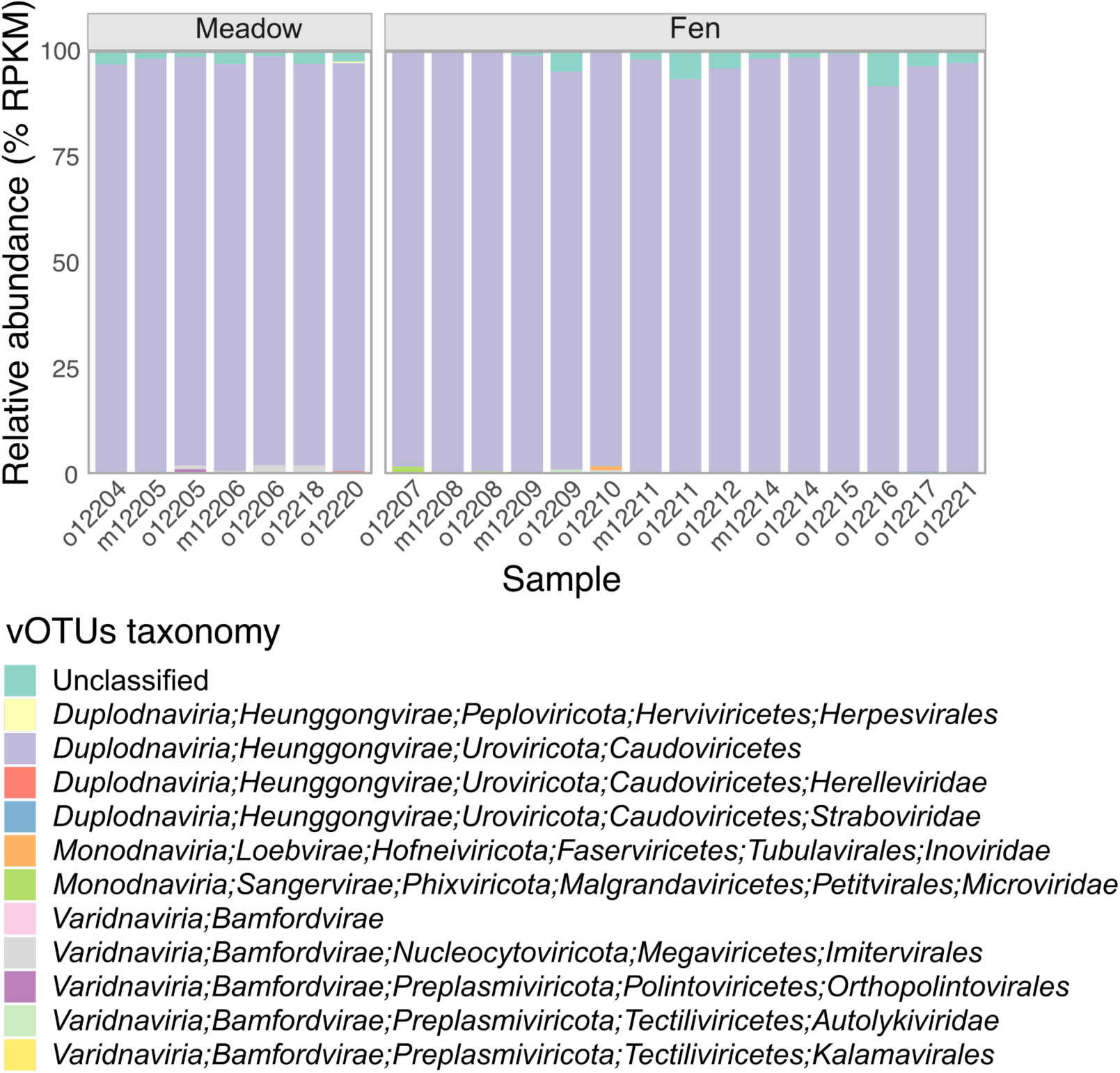
Relative abundance of different taxonomic groups assigned to Kilpisjärvi vOTUs.

**Figure S4.**
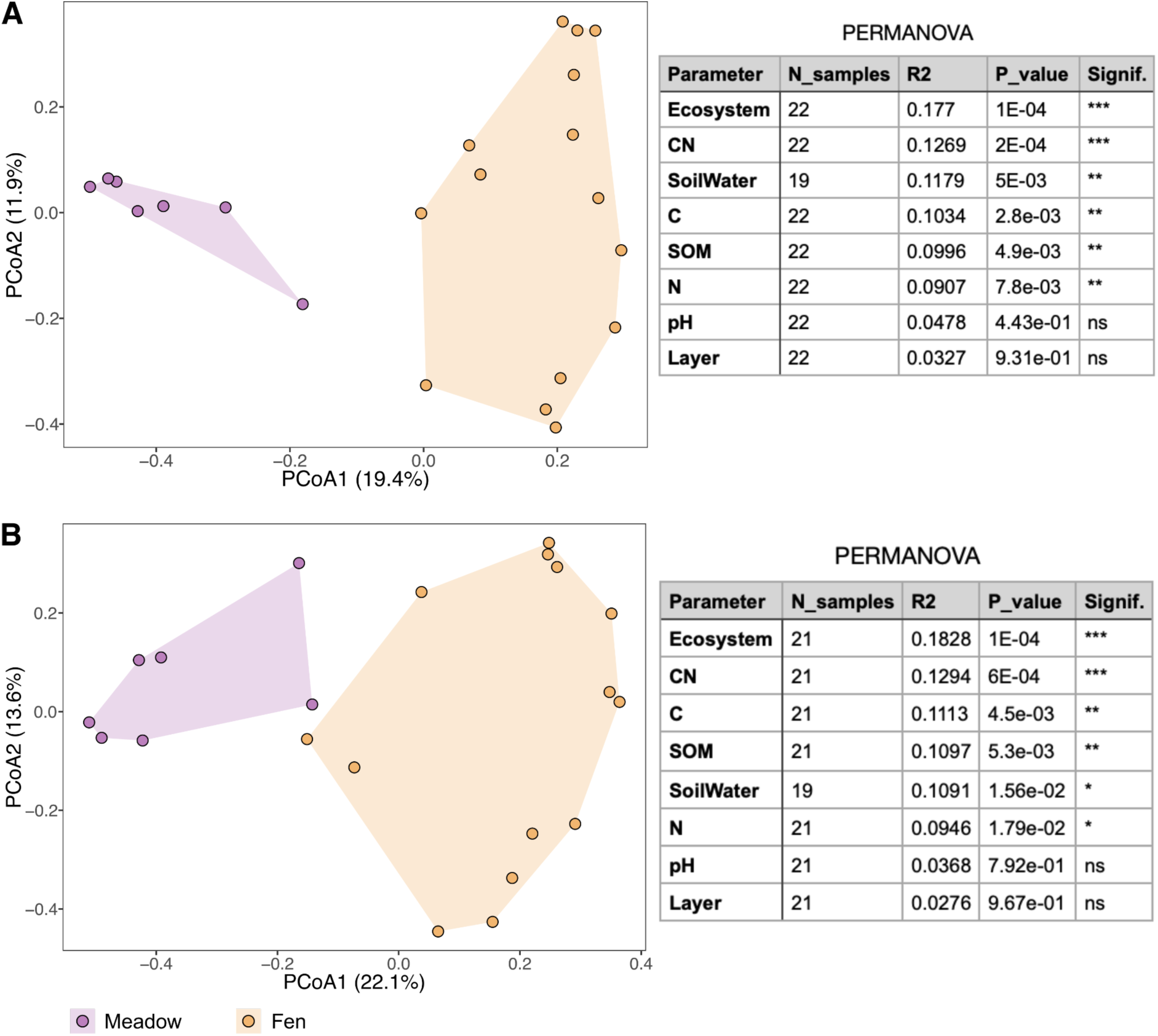
PCoA of (A) Kilpisjärvi vOTUs (n=1881) and (B) *Acidobacteriota*-associated vOTUs (n=125) in 22 Kilpisjärvi meadows and fens metagenomes. Convex hulls show actual spread of points. The minimum 50% horizontal coverage was applied. The R2 and p-values were obtained separately for each variable.

**Figure S5.**
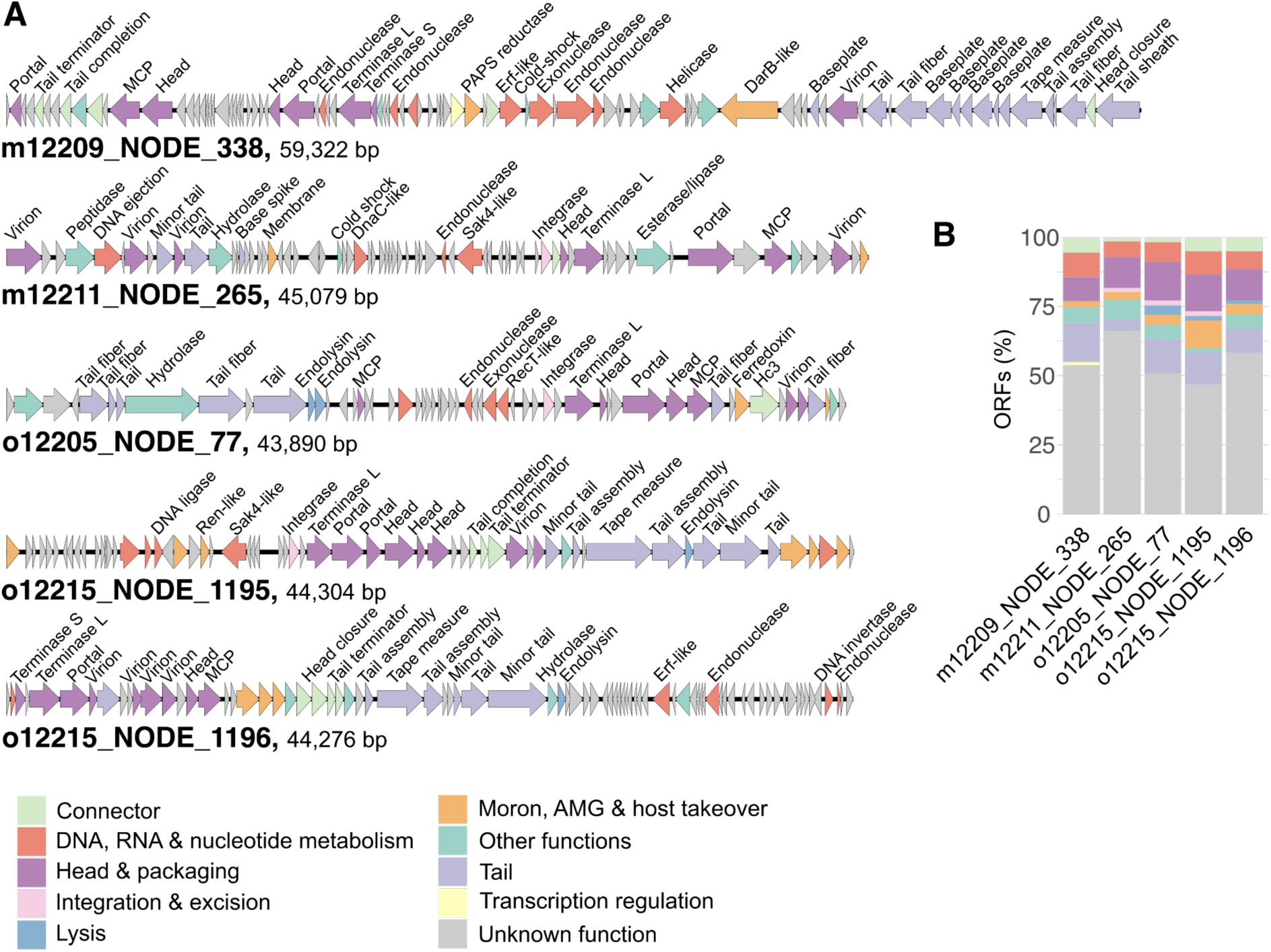
High-quality (96-100% complete) vOTUs assigned to *Acidobacteriota*. (A) Genomes with ORFs shown as arrows and colored according to the functional categories (see the color key). (B) Distribution of ORFs according to the functional categories, same color key as in (A).

**FigureS6.**
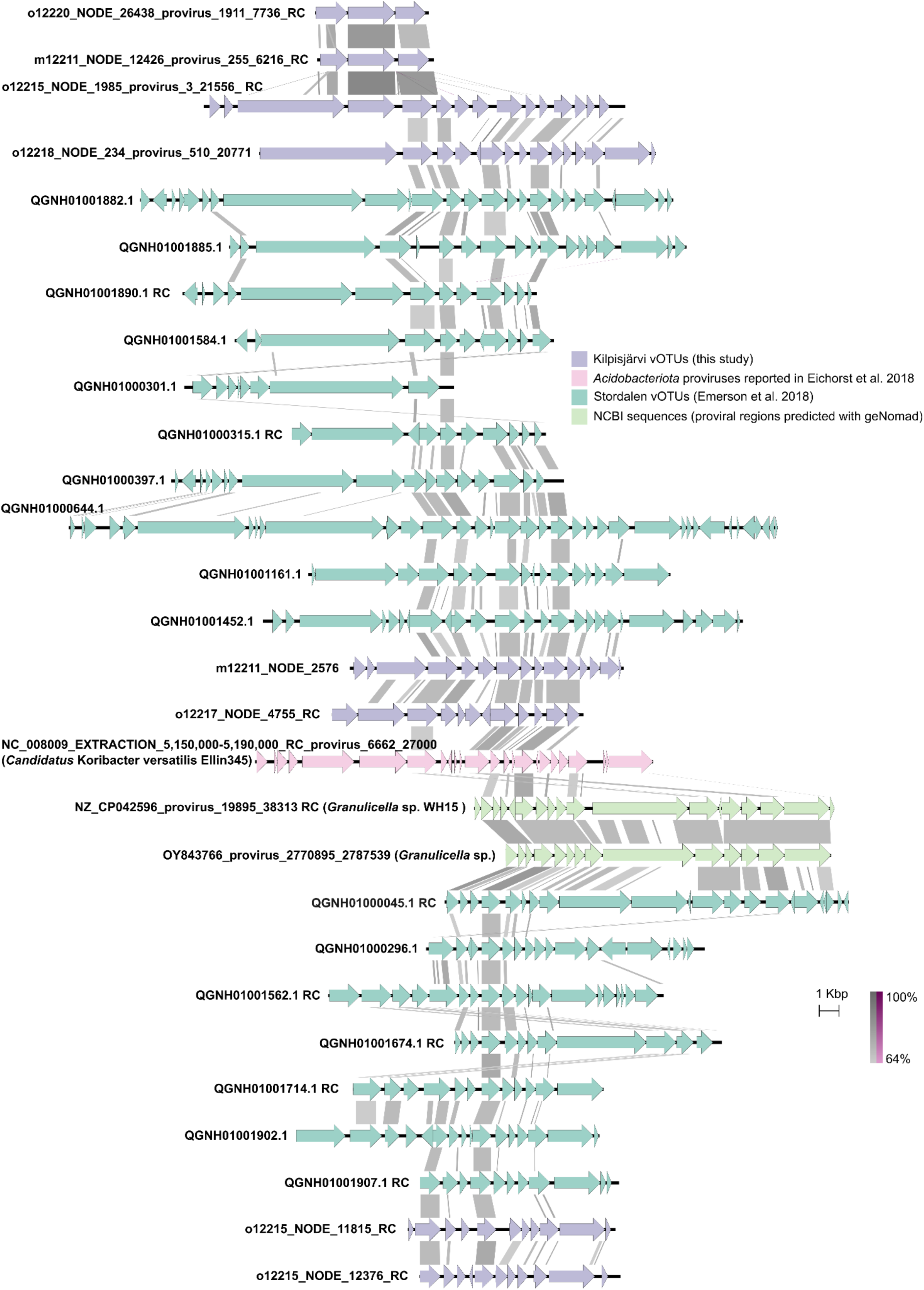
Putative acidobacterial proviruses identified in this study and related vOTUs reported in Emerson et al. 2018 (PMID: 30013236), *Candidatus* Koribacter versatilis provirus reported in Eichorst et al. 2018 (PMID: 29327410), and NCBI references. ORFs are shown as arrows and similarities between genomes (BLASTn) are in shades of gray (direct) or purple (invert). RC, reverse complement.

**Figure S7.**
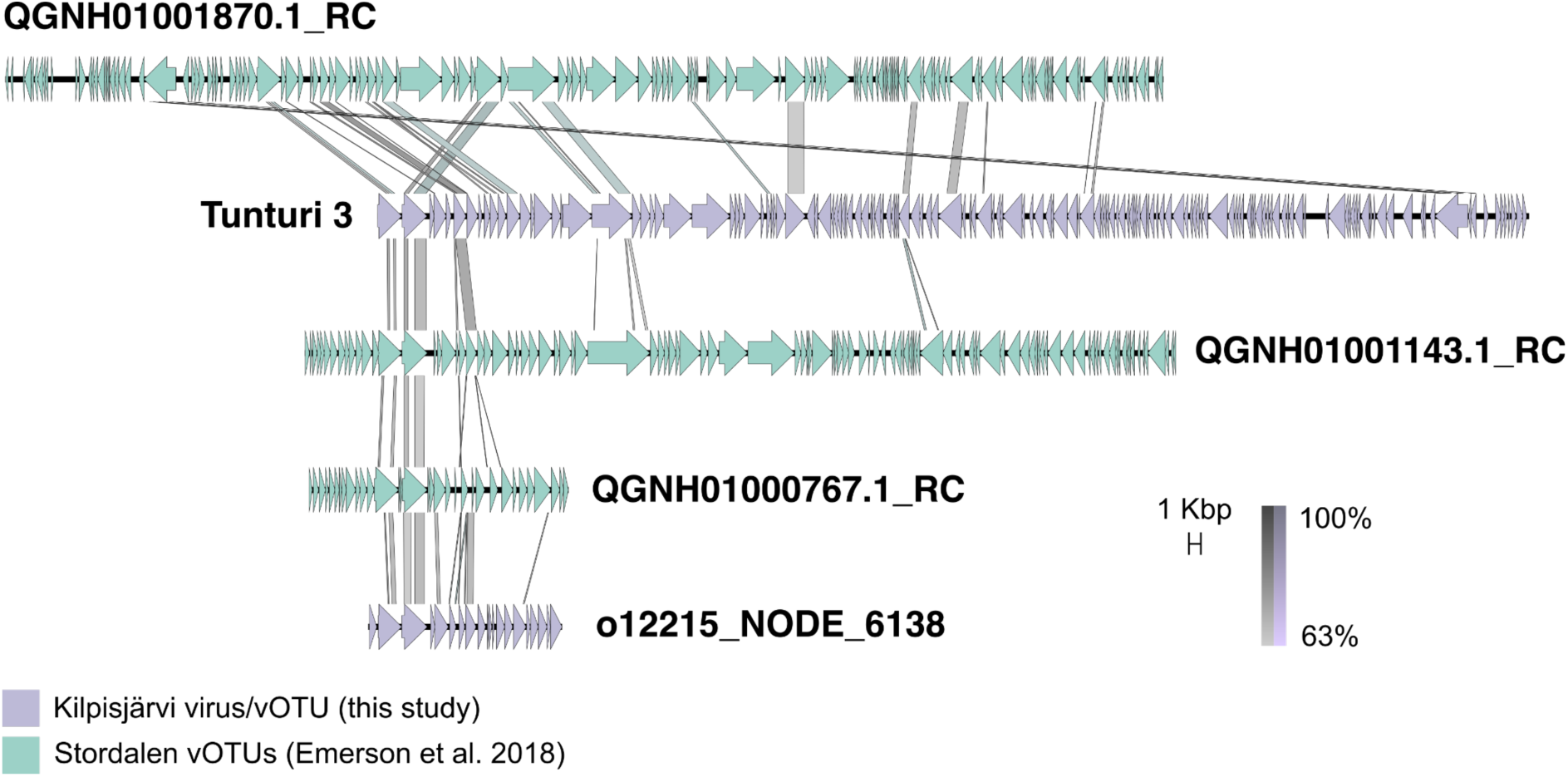
VConTACT2 viral cluster comprising the isolate Tunturi 3, o12215_NODE_6138 (this study) and three vOTUs reported in Emerson et al. 2018 (PMID: 30013236). Similarities between the genomes (BLASTn) are shown with the shades of gray (direct) or purple (invert).

